# Towards the definition and measurement of routines and the cognitive processes that underpin their maintenance

**DOI:** 10.64898/2026.03.26.714585

**Authors:** Christopher R. Nolan, Mike E. Le Pelley, Kelly G. Garner

## Abstract

The benefits of routines for daily functioning are widely acknowledged, yet, despite their apparent importance, methods for quantifying routine maintenance and the causes of their disruption remain lacking. Here, we propose a novel means of defining and quantifying routines (transition entropy). Using the transition entropy, we show that routines can be robustly elicited on tasks that require searching through a grid of squares for a hidden target. Over two experiments (N=100 each), we show that use of routines—as quantified by transition entropy—is robustly perturbed by frequent switches between search grids, as locations specific to the currently irrelevant grid become competitive for selection. Using a normative model that tracks task dynamics, we show that disruption to routines can be attributed to altered sensitivity to the odds of success for completing a task. This suggests that routine maintenance may be disrupted by over-sensitivity to a lack of reward early in routine performance, or increased expectations regarding the utility of pursuing other tasks.

---

It is easy to intuit that maintaining a good routine supports the task of getting up and out of the house. Despite the apparent importance of following a regular order (or set of orderings) in behaviours for health (Finucane et al., 2025; Mindell & Williamson, 2018; O’Conor et al., 2019) and productivity (Clear, 2018), our understanding of the cognitive processes that underpin routine maintenance remains under-explored. Indeed, theoretical and empirical investigations into how humans manage ordered behaviours typically prescribe a correct order of events (e.g. Carlson & Sohn, 2000; Garr & Delamater, 2019; Graybiel, 1998; Humphreys et al., 2000, 2001; Schneider & Logan, 2006; Watanabe et al., 2023; T. Wen et al., 2020; X. Wen et al., 2025). Although this allows for analytical simplification, such procedures more closely correspond to deterministic sequences that underpin automated chunks of behaviours, such as the expert performance of a piano concerto. With routines, however, what is deemed as the “correct” order of behaviours appears to be in-part determined by the individual — it is not necessary that one has breakfast before getting dressed. Until there exists a framework for the theoretical and experimental analysis of the self-ordering of task-relevant behaviours, we will lack a rigorous definition of routines, as well as an understanding of the factors that shape them, their maintenance, and their consequent benefits.

The tendency to assemble behaviours into a regularly (but not definitively) followed order to complete a recurring task is apparent even in the absence of explicit reward structures that favour such a strategy. Desrochers et al. (2010) demonstrated this by presenting macaques with grid displays made up of nine dots, where the task was to fixate the dots until landing on a randomly designated target. Despite the random allocation of the rewarded location, the macaques showed consistent orderings of fixated locations that were maintained over periods, that changed in configuration several times over the course of the experiment. The likelihood of making an adjustment was influenced by the extent to which the length of the most recent ordering exceeded the average sequence length (i.e., the average number of fixated locations used to attain a reward). This suggests that temporal cost may serve as a feedback signal that drives the ordering of behaviours (Desrochers et al., 2010, 2015), potentially by determining the average reward accrued per time step (Daw & Touretzky, 2000, 2002; Dezfouli & Balleine, 2012; Dezfouli et al., 2014; Mahadevan, 1996; Tsitsiklis & Van Roy, 2002). Thus, the tendency to form routines may exist in order to optimise the cost-benefit ratio for behaviours that must be performed over an extended period of time. Despite this apparent tendency to form routines and the implied benefit, we often fail to maintain existing routines despite being motivated to do so (Gardner et al., 2024; Rothman et al., 2015). This may seem surprising given the apparent negative consequences for daily functioning when routines fail (Arlinghaus & Johnston, 2019), and suggests that further investigations are required to uncover the sources of disruption to routine maintenance (de Wit et al., 2018).

One reason that routines may be challenging to maintain is that they occur in environments that support multiple known routines. For example, a host of recipes are prepared in the kitchen, each using common components such as the stovetop. There is a growing body of knowledge from the study of single behaviours—such as pressing a button to a stimulus—that suggests that such overlap should make it harder to select behaviours relevant for the intended routine (see Hommel, 2019, 2022; Kiesel et al., 2010, for reviews). Specifically, it has been shown that response times are slower when stimuli possess features relevant to the task performed on the previous trial, even when the response currently required is exactly the same as the previous response (Rogers & Monsell, 1995). This suggests that common sensory elements may serve to activate behaviours relevant to multiple routines, decreasing the probability that the current routine is maintained.

Whether such findings extend to ongoing tasks consisting of multiple ordered behaviours remains unexplored. To begin addressing this question, Garner et al. (2024) had human participants search a 16-square grid to find a hidden target, by making eye movements to reveal the contents of each square. The grid was bordered by one of two colours, where each colour signalled a unique task (Task *A* or Task *B*) comprising a non-overlapping subset of four locations. On each trial, the target location was chosen at random from these four task-relevant locations. The number of observed transitions between squares to find a target on each trial decreased over the course of the session, suggesting the development of routines that were specific to each task. Further, erroneous fixations towards Task-*B*-relevant locations were higher during performance of Task *A* (and vice versa), relative to fixations towards locations that never contained a target. Thus cross-task cues (in this case, possible target locations) exert more control over behaviour during performance of a routine, relative to irrelevant cues. What remains to be understood is what causes such cross-task cues to derail established routines; specifically, what are the factors that render routines more or less vulnerable to disruption from cross-task interference? Further, given the anecdotal prevalence of failures in daily routine maintenance, we ask whether there are potential benefits to be gained from reduced routine adherence, and do these benefits help us understand why routines might be susceptible to disruption?

One factor that may modulate the extent to which cross-task cues disrupt routines is the extent to which routines can be mentally separated from one another despite their occurrence in common environments. Performing an ongoing task (such as completing a routine) requires selective representation of the rules that link stimuli and responses to desired outcomes (e.g. Niv, 2019). The extent to which we can mentally separate tasks that occur in overlapping environments appears to rely on the extent to which transitions between tasks co-occurs with distinctive events (Flesch et al., 2018, 2022; Gershman & Niv, 2013; Gershman et al., 2013, 2014). The continued application of outdated task-rules (i.e. the failure to mentally separate a previous task from the current task) reduces when the change in task-context coincides with an abrupt change in stimulus-outcome relationships (Gershman et al., 2013), a larger difference between task-relevant sensory-inputs (Gershman et al., 2014), or when transitions between tasks are rare compared to more frequent (Flesch et al., 2018). Collectively, these findings suggest that encountering an event that is relatively surprising signals that the world has changed, and that a new set of response rules can be applied in the same environment. Relatedly, an assumption of theoretical accounts seeking to explain how cross-talk influences performance of stimulus-response rules is that between-task interference decreases when there is greater separation between task representations (see Feng et al., 2014; Garner & Dux, 2022, 2023; Musslick & Cohen, 2021, for reviews). This suggests that cross-talk, and by proxy, routine maintenance, should be modulated by the extent to which switches between tasks are surprising, as surprising switches enables stronger mental separation of the response-rules that govern each task. However, these theories also assume a cost to mentally segregating task representations to reduce cross-talk; the more separated a task-representation, the less the contents can be shared to support performance on novel but related tasks. This suggests a possible benefit to keeping routines in a state of vulnerability to cross-talk; reduced mental separation of routines may serve to increase options when reconfiguring behaviour for novel tasks.

A recent study provides the conditions for answering the question of whether the rate of switching between tasks serves to mentally segregate representations in the service of routine maintenance. Using an adaptation of the grid-search paradigm described above, Barnes et al. (2026) determined the impact of rare vs frequent task transitions on the generalisation of learned behaviours to new tasks. Once again, two coloured borders signalled which unique subset of four locations within the grid should be searched to find a target, this time by clicking on squares using a mouse to reveal their contents. After establishing accurate performance on each task, participants completed a rehearsal phase where they performed both tasks under conditions of a rare (5%, Stable group) or more frequent (30%, Variable group) probability of a task-switch during which time the coloured borders were removed. The authors tested whether switching during rehearsal impacted performance on a novel task with a new coloured border where the key locations were pseudorandomly drawn from each of the trained tasks (e.g. *A*_1_, *A*_2_, *B*_1_, *B*_2_: mixed transfer), relative to when a new border signalled a full replication of one of the trained tasks (e.g. *A*_1_, *A*_2_, *A*_3_, *A*_4_: identity transfer). As anticipated, the frequency of task-switches during rehearsal affected the rate of acquisition of mixed transfer relative to identity transfer; participants in the Stable group—who had less experience in shifting between tasks—showed a greater difference between transfer tasks (with mixed transfer performance being poorer than identity transfer performance) than did participants in the Variable group. This result is consistent with the idea that rare switching during the rehearsal phase served to better separate the two task representations, rendering it more difficult to make associations between them to solve the mixed transfer task (cf. Lee et al., 2022). However, the authors did not determine whether the task-switch rate during rehearsal impacted the strength of routine maintenance, nor did they determine whether routine strength during rehearsal was detrimental to learning-transfer performance. Addressing these questions will help determine whether performing a task under more volatile conditions is indeed a source of routine disruption, and whether disrupting routines may be desirable under some circumstances, such as through increasing flexibility under conditions of learning transfer.

As mentioned above, Barnes et al. (2026) removed the coloured borders during rehearsal, where these colours had previously signalled whether Task *A* or *B* should be performed. Under these conditions, finding the target in the fewest moves on average requires exhausting the locations from the task that yielded a target on the previous trial, before moving to the other task (this is true for as long as the probability of a task switch is less than 50%). Over two experiments, the authors found that instead of using this strategy, the Variable group performed higher numbers of pre-emptive location selections from the less-likely task (e.g. Task *B*), before exhausting the more-likely task (e.g. Task *A*), compared to the Stable group. This further suggests that more frequent switching resulted in poorer separation of the two task representations, but it remains unknown whether frequent switching impacted routine maintenance. The observed behaviour could be produced by disruptions to routine maintenance, or by the assimilation of cross-task behaviours into an ongoing but apparently suboptimal routine. Adjudicating between these possibilities allows us to not only understand the mechanisms via which routines are made more or less effective, but also into the processes that drive flexible performance on novel tasks. However, there is currently no ideal metric for quantifying how consistently a particular order of location selections was followed, the absence of which is a major limitation when determining the causes of weakened routine maintenance.

Some existing metrics quantify elements of routine-like behaviours. Previous work examining how macaques self-order fixations has primarily focused on quantifying the trial-by-trial cost of orderings relative to an average (Desrochers et al., 2010, 2015), or the energetic factors driving the formation of routines over time (Brynildsen et al., 2026), rather than quantifying how regularly routines have been followed. The latter is required when seeking to order individuals according to how strongly they maintain a routine. Garner et al. (2024) quantified routine maintenance by calculating the variance of an individual’s transition probability matrix, where the matrix codes how frequently there was a transition between each pairing of grid locations. This measure goes some way to indicating variance over performed orderings, which can serve as a proxy for the extent of routine maintenance. For example, the variance of the collective transition probabilities of an individual who more frequently performed two of four possible transitions (e.g. .4, .4, .1, .1) will be higher (*s*^2^ = 0.03) than the variance when transitions were performed without any preferred order (e.g. .25, .25, .25, .25, *s*^2^ = 0). However, this metric is potentially confounded with task accuracy: variance of the transition probability matrix also changes with the total number of transitions used. Routines that include extraneous behaviours will result in a higher variance score relative to those that do not, even if comparably consistent ordering occurs on every trial. For example, if only 2 of the transitions are strictly required to perform the task, the participant who learns this will use fewer transitions (e.g. .5, .5, .0, .0) and consequently show higher variance (*s*^2^ = .083), than one who has mistakenly added a third transition, but performs each transition as consistently (e.g. .33, .33, .33, 0, *s*^2^ = 0.027). Thus, an approach is required that can quantify the regularity of behaviour that is not contaminated by task accuracy (i.e. the number of locations visited while performing the task).

Here, we present a novel approach that overcomes this limitation in quantifying the extent to which a routine has been exhibited. We present a computation of the entropy over transitions that ranks people according to how variably they leave locations (or states), on average, while being agnostic to the total number of locations (states) visited (here termed the *transition entropy*). This metric unlocks the ability to quantify regularity in the implementation of any set of events or behaviours, thereby opening up the experimental investigation of the costs and benefits of routine maintenance. Here, we seek to tease apart whether cross-task behaviours disrupt routines, or become assimilated into an entrenched routine. We determine, via a new analysis of the data from Barnes et al. (2026), whether the rate of switching between tasks systematically modulated routines during rehearsal as indexed by transition entropy. If volatile environments challenge routine maintenance, then the Variable group should show higher transition entropy than the Stable group. In contrast, if volatile environments induce stricter maintenance of inefficient routines, then the Variable group should show lower transition entropy. As this switching manipulation was present in the rehearsal phase of two experiments in Barnes et al. (2026), we were also able to determine whether this effect replicated across experiments.

Upon establishing the impact of switching on routine maintenance, we next sought to determine the causes of failures in routine maintenance. If routine maintenance is weakened due to decreased surprise when switching between tasks, then poorer routine maintenance should be associated with altered sensitivity to the probabilities that underpin computation of the associated likelihood. Testing this hypothesis, using a normative model that tracks probabilities of the tasks’ dynamics, was the second key aim of the current study. Finally, we sought to determine whether routine maintenance, which may depend on the mental separation of tasks, is achieved at the expense of generalising knowledge to new tasks, which depends on reduced separation between representations. We therefore tested whether stronger routine maintenance predicted poorer learning transfer scores as observed by Barnes et al. (2026). If routines are more easily maintained under conditions of greater discernibility between task representations, then reduced transition entropy should correspond to poorer performance on a novel task that requires a novel combination of routine elements (mixed transfer) relative to a task that requires replication of a routine (identity transfer).

## Results

The first key aim was to derive a measure that indexes how strongly a routine has been maintained during task performance. Participants were presented with a grid array of 16 squares, and learned to click on squares using a mouse button until they found an animal target image hidden behind one of the squares (Fig 1A). In a task acquisition phase, the grid was bordered by one of two colours, where each colour signalled which unique subset of four locations within the grid should be searched to find the target (Fig 1B). These two ‘tasks’ are hereon referred to as Tasks *A* and *B*. Participants were instructed to learn to find the targets in four moves or fewer.

**Figure 1.**
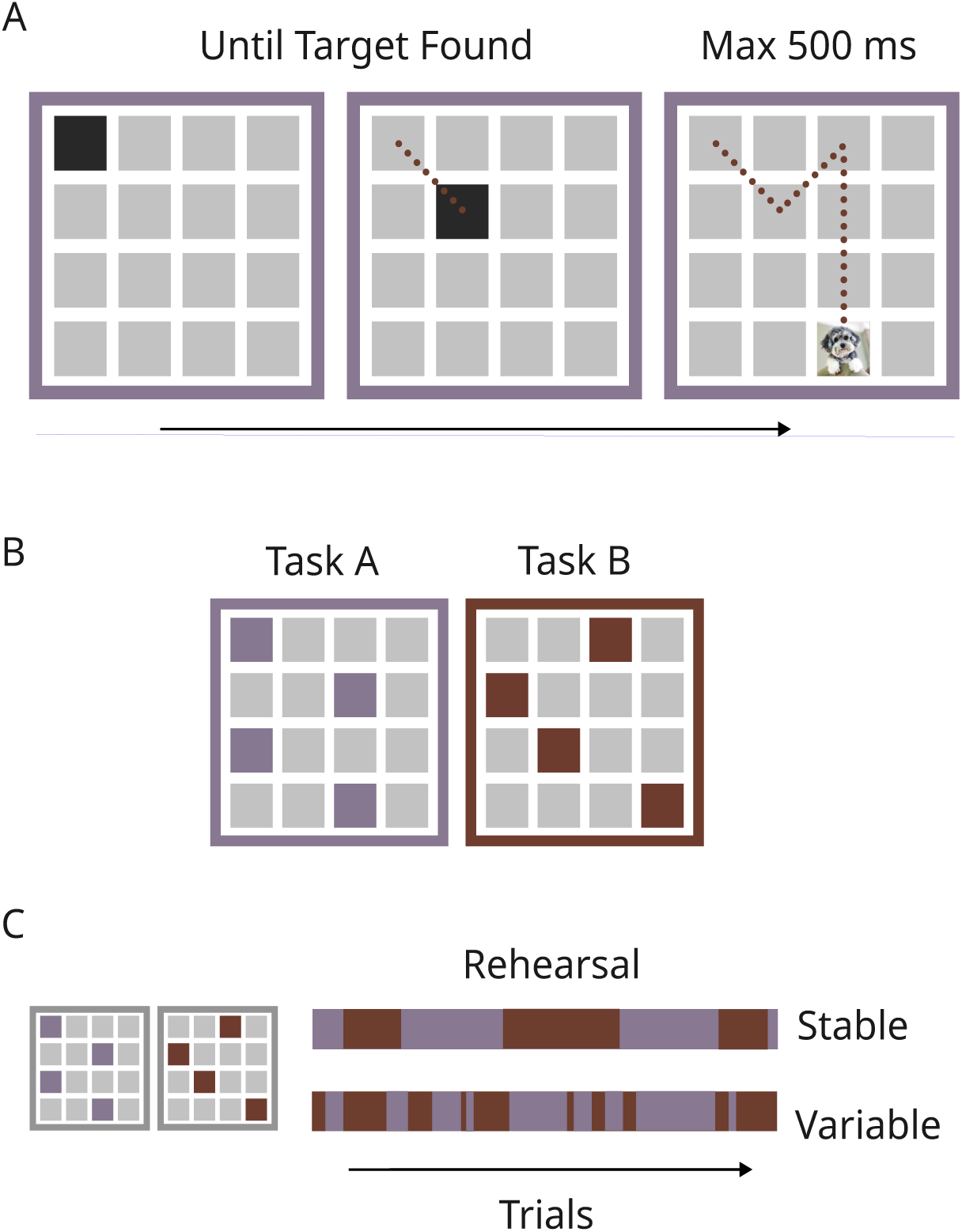
Experimental task. A) Example of a trial: participants clicked on squares using a mouse. Upon clicking, the square either turned black to signal no target, or the target image was shown. The target was shown for a maximum of 500 ms, including the time for which the mouse button was pressed. B) Task A and Task B: depending on the colour of the border, the target would be found at only four locations. On each trial, the target was randomly allocated to one of the four task-relevant locations. C) Schematic of the rehearsal phase from Experiments One & Two: During rehearsal, the coloured borders were replaced with a single grey one. Participants were allocated either to the Stable group, with a probability (p)=.05 of a task-switch, or to the Variable group, with p(task-switch)=.3.

After task acquisition, participants completed a rehearsal stage; the goal of the rehearsal stage was to manipulate expectations regarding the likelihood of upcoming cross-task events. Participants were informed that their task was to continue to find the targets in four moves or fewer. Further, they were informed that they would no longer see the coloured borders, and that now the task could switch on any trial. Participants were also informed that the task was always more likely to stay the same (Stay trials) than to switch (Switch trials). Note that if the probability of a switch is less than p=.5 then finding the target in the fewest moves possible on average requires exhausting the task locations from where the last target was found before moving to the other task. Participants were randomly assigned to one of two groups. For the Stable group, the probability of a task switch was p=.05; for the Variable group the probability was p=.3 (Fig 1C).

### A Measure to Index Routine Maintenance

Previous analysis of the rehearsal phase data showed that the Variable group were more likely to make pre-emptive visits to locations relevant for the less-recent task before exhausting locations from the most-recent task (Barnes et al., 2026). The Stable group, having just found a target from Task *A*, were more likely to (optimally) exhaust all Task *A* locations before moving to Task *B* locations if necessary. An example of behaviours from two participants is presented in Fig 2D). Note that both groups showed low and comparable numbers of location selections that were irrelevant to both tasks. An index of exhibited routine is required to determine whether pre-emptive visits to the less-recent task (exhibited by the Variable group) reflect a disruption to routines, or a routine that includes cross-task behaviours.

**Figure 2.**
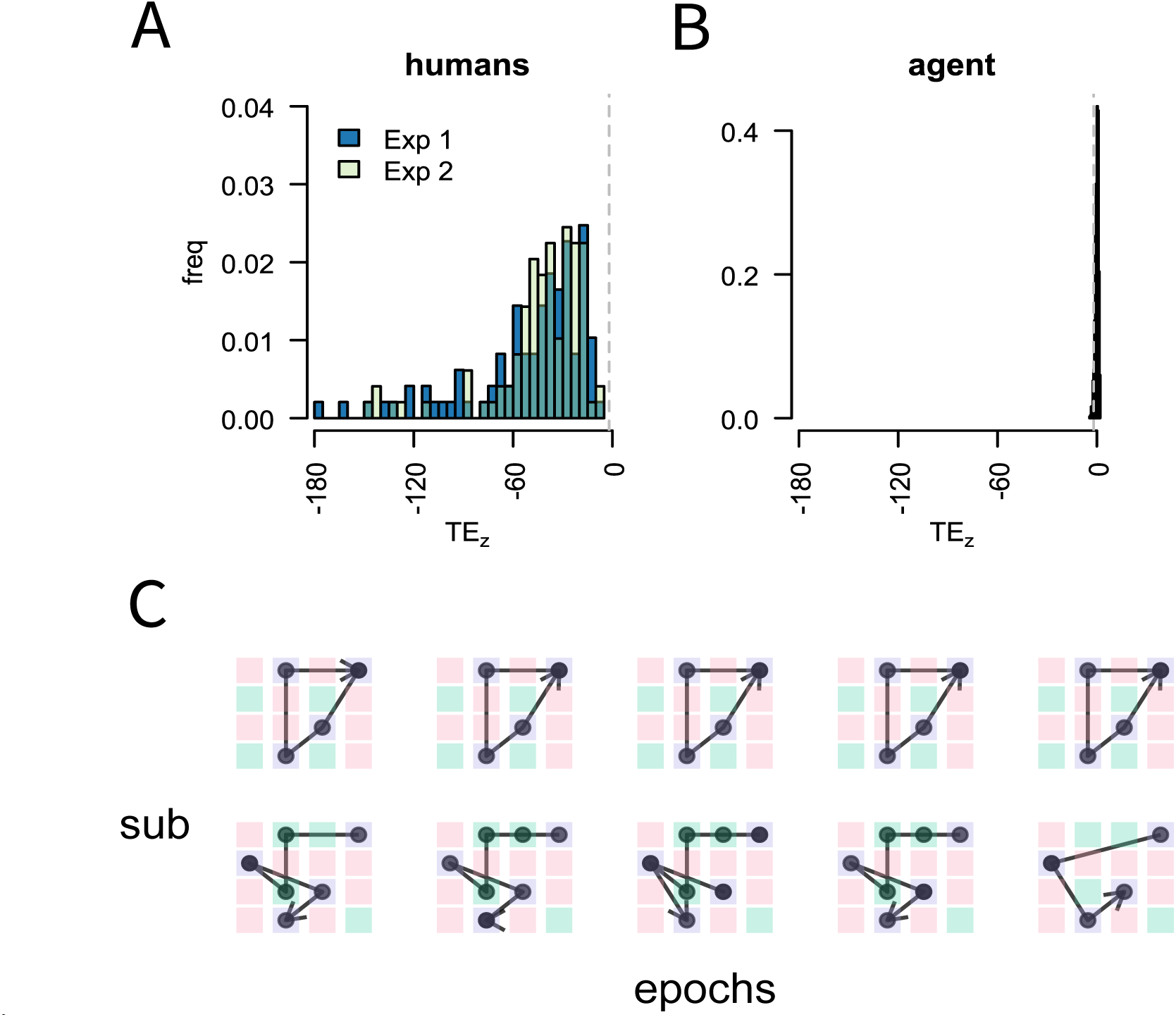
TE scores are different from chance. A) Left histogram is of TE_z_ scores from Experiment One (Exp1) and Experiment Two (Exp2) and the right histogram (B) is of TE_z_ scores for an agent that performs accurately but orders responses randomly. Vertical dashed lines indicate z=-1.96. C) Example location selections from two participants (rows) over epochs (columns) selected to show a complete visit to all task relevant locations (blue squares). Green squares show locations relevant to the other task, and red squares never contained a target. freq = frequency, sub = subject

#### Transition Entropy [TE]

To quantify how strongly individuals followed particular orderings of location selections over the long run (i.e., the extent to which they maintained a routine), all location transitions made in a given task state (*A* or *B*) over the course of the rehearsal phase were summed into a counts matrix, where rows reflect the location (state, *s*) that is being moved from, and columns reflect the location (state, *s^′^*) that is being moved to. Each row of the transition counts matrix was normalised into probabilities, so that each row coded the probability of each *s^′^*, given an *s*. To calculate how regularly a state was departed, entropy scores were calculated for each row of the matrix, using the formula:

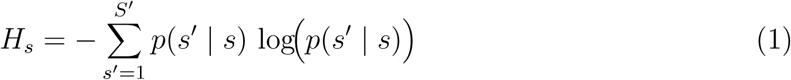

where *H_s_* is the entropy for a given state *s*. Higher entropy values correspond to greater disorder in the ordering of behaviours when leaving state *s*.To aggregate the state entropies into a single score, a weighted sum was performed, where each *H_s_* value was weighted by the probability of a visit to that state.

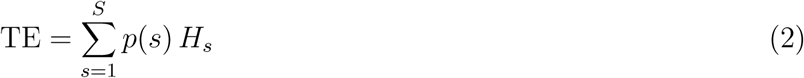

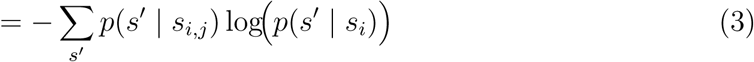

where *j* counts over the columns of the transition matrix. Note that this formulation is very similar to the Shannon entropy, except that the logged probability is conditioned by the state rather than by all observations. This score thus reflects the expected variation when leaving a state. As the score is based on the number of ways states are traversed and not the number of states visited, location selections that follow comparable ordering strength will receive the same TE score that is robust against differences in the number of elements (states) that make up the routine. For example, the location selections *A*_1_ *→ A*_2_ *→ A*_1_ *→ A*_2_ result in the same TE score as *A*_1_ *→ A*_2_ *→ A*_3_ *→ A*_1_ *→ A*_2_ *→ A*_3_, as each *s^′^* is always transitioned to from the same *s* (and this applies to the probabilistic as well as the deterministic case). Further, the measure scales according to how many transition types are made. For example, the TE score is the same when a single *s^′^* is entered from four different *s*, or two *s^′^* are each entered from two different *s* (given comparable transition probabilities across the two scenarios). Note that a greater TE score corresponds to greater variation in the ordering of responses; thus a higher TE score suggests poorer maintenance of routines over the long run. This formulation of TE provides a novel definition of what it means to adhere to a routine. Viewed through this lens, the more predictable the expectation regarding the next state, the more a routine has been maintained.

### Determining Elicitation of Routines

Given this novel means to index routine maintenance, the next goal was to determine whether the rehearsal phase robustly elicited routines, i.e. a regularity in the order of location selections, that occurred to an extent that was greater than what would be expected by chance. To achieve this, we permuted a null distribution of TE scores for each participant, based on random selections from observed responses over 1000 iterations. We then z-scored the observed TE score for each participant (TE*_z_*), using the mean and standard deviation of their permuted null TE scores. We reasoned that as lower TE scores reflect greater maintenance of a routine, then TE*_z_* scores should be lower on average than -1.96 (i.e. be likely to be observed less than .025% of the time under the null). A one-sample t-test confirmed that TE*_z_* scores were statistically lower than -1.96 both for Experiment One (M=-50.29, SE=3.67, t(96)=-13.17, p<.001), and for Experiment Two (M=-43.79, SE=2.93, t(97)=-14.28, p<.001, Fig 2A). Thus, participants maintained consistent orderings of location selections that occurred to an extent that was greater than chance.

Next, to benchmark observed TE*_z_* scores against accurate but random performance, a perfectly performing, randomly ordered agent was used to generate a selection of task-relevant locations (e.g. selecting only *A_i_* locations, and not *B_i_* locations, while performing Task *A*) in order to complete the rehearsal phase of the experiment, with the constraint that the same location could not be selected twice in a row. Given this set of randomly ordered responses, a null distribution was generated using the same procedure as described above, and the consequent null TE*_z_* was calculated. This process was repeated over 1000 iterations, to generate a distribution of TE*_z_* scores for the agent. We then took the mean TE*_z_* scores (reported above), and determined the likelihood of an observation at least that small given the distribution of TE*_z_* scores derived using the agent (M=0, SD=0.98, Fig 2B). The probability of the observed TE*_z_* scores was p<.001 for both Experiment One and Experiment Two.

### More Frequent Switching Increases TE Scores

Having established that the paradigm elicits regularly ordered behaviours, we next investigated how the rate of switching between tasks impacted routine maintenance. In line with the idea that a volatile environment impairs routine maintenance, TE scores in Experiment One were statistically higher for the Variable group (M=15.06, SE=1.25) compared to the Stable group (M=9.81, SE=0.95, t(90.68)=2.37, d=0.47, 95% CI[0.11, 0.84], p=0.02, Fig 3A). This result is robust, as it replicated across experiments (Experiment Two: Variable M=17.7, SE=1.37, Stable M=11.57, SE=1.04, t(90.96)=2.6, d=0.52, 95% CI[0.15, 0.86], p<.001, Fig 3B). This between-group TE difference shows that the increased pre-emptive cross-task location selections shown by the Variable group in Barnes et al. (2026) can not be attributed to the assimilation of cross-task behaviours into an ongoing routine. Thus, it is likely that more frequent switching disrupted routine maintenance.

**Figure 3.**
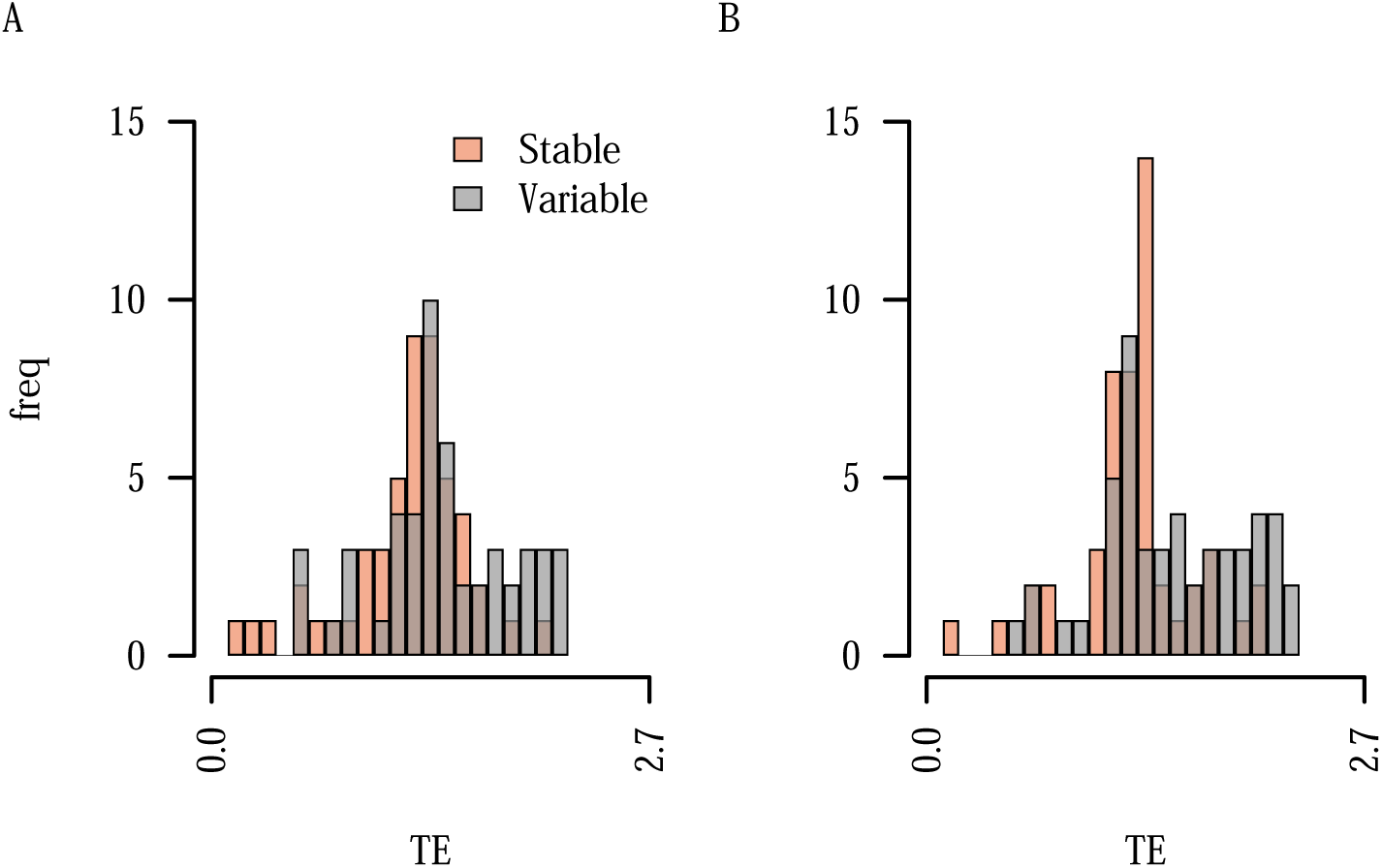
TE scores are reliably increased for the Variable group during Rehearsal. A) Histogram of untransformed TE scores from Experiment One for the Stable and Variable groups. B) Same, for Experiment Two. freq = frequency

### Cross-Task Interference Drives Disruptions to Routines

Having determined that more frequent switching disrupts routine maintenance, we next sought to explore why the Variable group exhibited increased cross-task selections during pursuit of a single target. Specifically, we asked whether these selections were due to altered sensitivity regarding the relevance of cross-task events. Given the removal of coloured borders in the rehearsal phase, finding the target in the fewest moves possible over the long run required exhausting the locations from the task where the last target had been found (Task*_LT_*) before switching to the other task (Task*_¬LT_*). Altered sensitivity that covaries with the odds that Task*_LT_* remains the current task (see Equation 6, *Methods*), or with the odds of success when sticking with Task*_LT_* (Equation 9, *Methods*) could motivate a performance strategy that does not match the normative strategy. Thus, we sought to determine whether increases in cross-task location selections covaried with variation in either the task-odds or the odds of success. Note that in so doing, we do not assume that people are Bayesian reasoners. Rather, we seek to determine how and where behaviour deviates from a normative benchmark. This provides clues for generating further hypotheses regarding how and why people may respond as they do to the task environment. To test this, we calculated trial by trial estimates of the task-odds and the odds of success, based on each participant’s history of location selections and target locations up to that trial, using a Bayesian normative model (note that the task odds were also updated during each trial, based on the location selections that occurred during the trial). See *Methods* for the details of this model. For each participant, the resulting values were entered into a hierarchical logistic regression where the dependent variable was whether a given location selection was from Task*_LT_* (0) or from Task*_¬LT_* (1). We also controlled for the general increase in experienced task-switches in the Variable group, by computing regressors to count the cumulative number of switches (Sw = Ʃ*_t_*_=1_*^T^S*(*t*), where *S*(*t*) = 1 if the task switched on trial *t*).

Across both experiments, the models showed good fit to the data (Figs 4A & 5A). Further, both models showed that the Variable group responded differently to variation in the odds of success, relative to the Stable group. In Experiment One, there was a significant group x task odds x success odds interaction (*B* = -0.06, *SE* = 0.02, LRT M1/M0 = 4.02, null M1/M0 *M* =0.41, *SD*=1.03, *p*=0.01). Specifically, the Variable group were most likely to make cross-task selections when the odds of success were lower, under conditions where the task odds were higher (see Fig 4B). Note that the odds of success remained constant over the course of a trial, and favoured continuing with Task*_LT_* as long as the probability of a task switch was less than 50%. The strength to which the success odds favoured Task*_LT_* was dependent on the probability of a task switch, with weaker favour conferred when the probability was higher. Note that although the probability of a task switch also impacted the task odds, the greatest changes in task odds occurred over the course of a trial, diminishing with every selection from Task*_LT_* that failed to yield a target. The current data showed that the Variable group’s differing response to the odds of success was evident when the task odds were higher, and diminished as the task odds decreased (see Fig 4B). Thus, the altered sensitivity to the odds of success exerted its influence earlier rather than later in the trial. The Variable group also showed an altered response to the odds of success in Experiment Two, as evidenced by a significant group x success odds interaction (B = -0.32, SE = 0.09, LRT M1/M0 = 3.42, null M1/M0 M=-0.98, SD=1.9, *p*=0.01, see Fig 5B). However, the group x task odds x success odds interaction was not statistically significant (p=.36). Overall, results from both experiments showed a group difference in the response to varying success odds, and suggest that variation in task odds might impact the response to the success odds.

**Figure 4.**
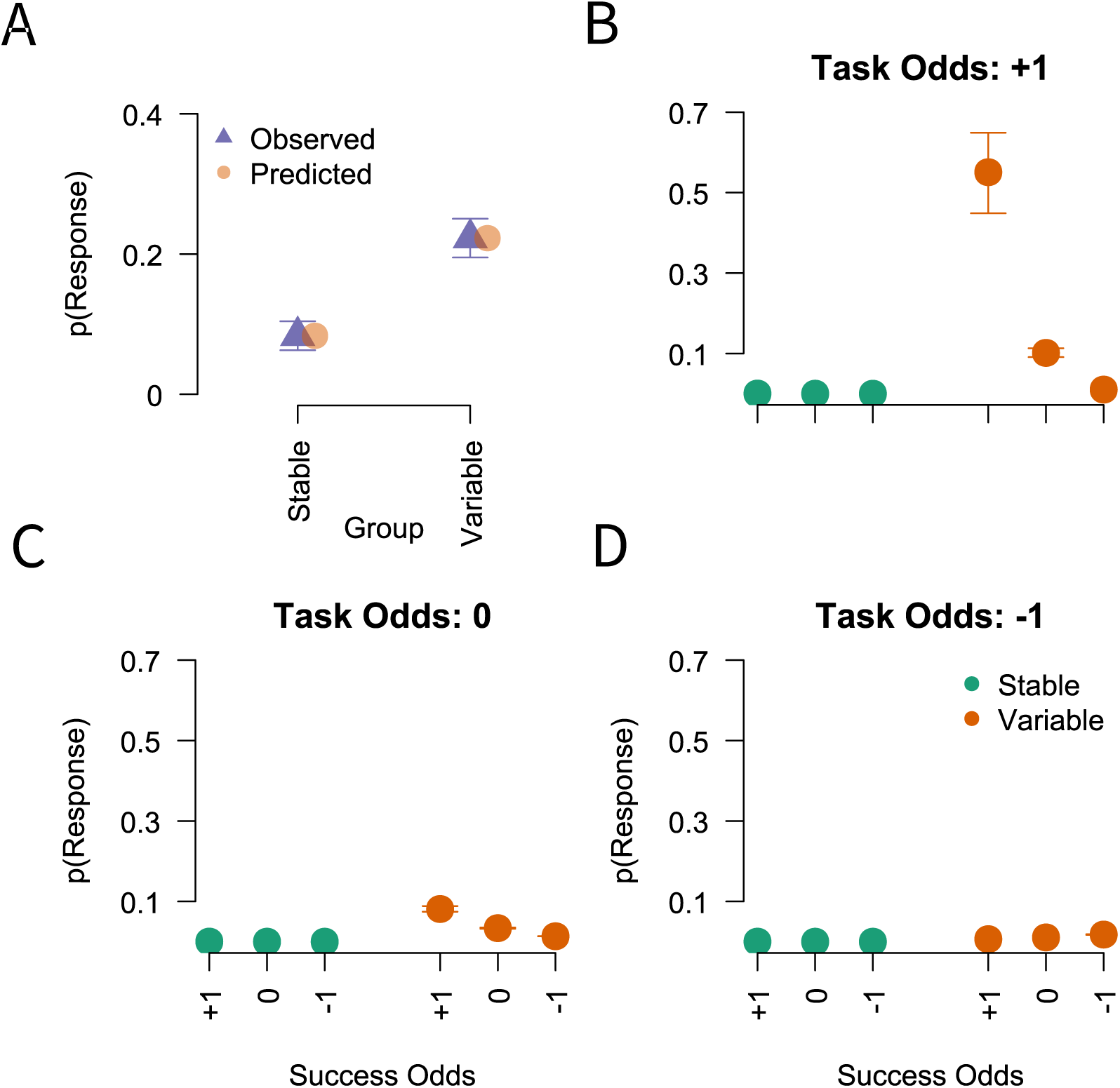
The influence of Group, success odds, and task odds) on cross-task selections in Experiment One. A) Observed and predicted probability of a cross-task selection (p(Response)), plotted by Group. B) Model predictions of the interaction between success odds (x-axis) and Group (coloured dots) when task odds was +1 SD above the scaled mean. C & D) Same as B, but for the mean scaled task odds (0) and for when the task odds was -1 SD below the scaled mean (-1 = -1 SD, 0 = scaled mean, +1 = +1 SD).

**Figure 5.**
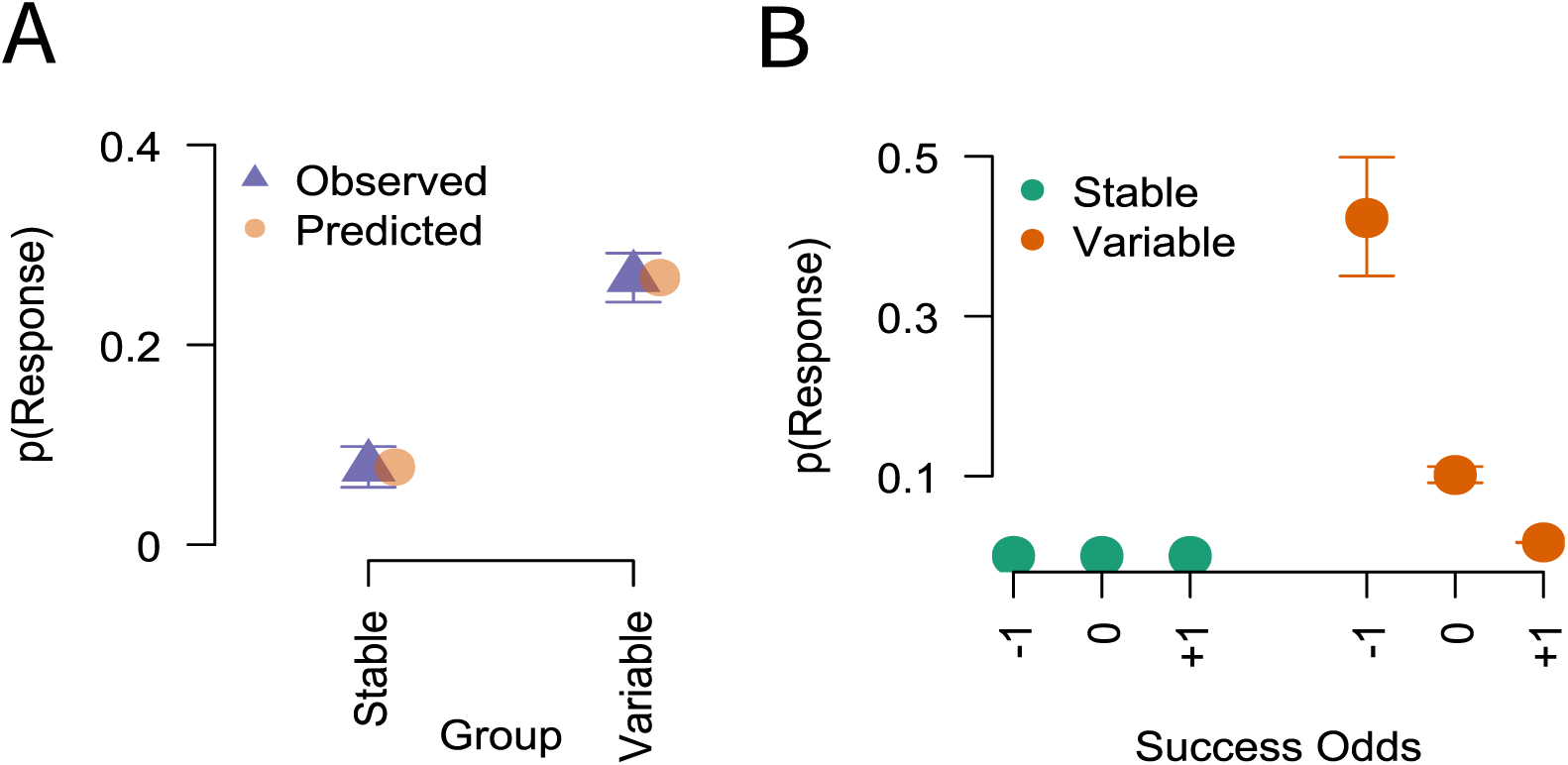
The influence of Group and success odds (-1 SD, M, +1 SD) on cross-task selections in Experiment Two. A) Observed and predicted probability of a cross-task selection (p(Response)), plotted by Group. B) Model predictions of the interaction between success odds (x-axis) and Group (coloured dots, -1 = -1 SD, 0 = scaled mean, +1 = +1 SD).

For each experiment, there was also a significant main effect of group (Exp 1: *B* = 6.08, *SE* = 1.1, LRT M1/M0 = 20.51, null M1/M0 *M* = -18.85, *SD*=6.05, *p < .*0001, Exp 2: *B* = 7.28, *SE* = 1.05, LRT M1/M0 = 28.83, null M1/M0 *M* =-20.99, *SD*=8.65, *p < .*0001), meaning that group differences remained even after accounting for variation in task odds, success odds, and the experienced number of switches. We return to this point in the Discussion.

### TE Scores are Associated with Learning Transfer Impairments

Having determined that routine maintenance is disrupted in the presence of increased cross-task selections, and that the frequency of cross-task selections is sensitive to the odds of success when sticking with a task, we next conducted an exploratory analysis to see if there are cognitive consequences of routine maintenance as indexed by TE scores. Specifically, if stronger routine maintenance is driven by higher expectations of success when sticking with a single task, then those who strongly maintain a routine might perform more poorly when required to combine behaviour subsets from across tasks (e.g. *A*_1_, *A*_2_, *B*_1_, *B*_2_, mixed transfer), relative to when replicating a known routine in the presence of a new coloured cue (e.g. *A*_1_, *A*_2_, *A*_3_, *A*_4_, identity transfer). Thus, there could be a negative relationship between TE scores and the time taken to learn the identity transfer task relative to the mixed transfer task.

To determine whether TE scores (which index the strength of routine maintenance) were associated with knowledge for mixed relative to identity transfer, transfer bias scores were calculated as the accuracy difference between the two tasks, where accuracy is calculated as the proportion of location selections that are task relevant, averaged across trials. Positive bias scores reflect greater accuracy for identity relative to mixed transfer, whereas negative scores reflect the opposite. In support of the idea that stronger routine maintenance corresponds to a reduced tendency to mix routine subsets, there was a statistically significant negative correlation between TE scores and transfer bias scores (*r*(87)=-0.33, 95% CI[-0.5, -0.13], *p*<.001, Fig 7). Note that Barnes et al. (2026) also estimated the ‘learning onset’—i.e., the point at which full knowledge of the transfer task had been gained—using an established Bayesian algorithm that evaluates ongoing performance as showing evidence for or against task knowledge (Maggi et al., 2024). We computed a comparable bias measure using this onset score, but did not observe a statistically significant relationship with TE scores (*p*=0.21). Thus the observed relationship between TE scores and transfer performance is dependent on the measure of performance used to compute the bias scores. We return to this point in the Discussion.

## Discussion

We sought to determine whether volatile environments impact routine maintenance, and whether interference from other tasks plays a role in observed impacts. To achieve this, we proposed a novel method for quantifying the regularity of ordered behaviours (TE, *transition entropy*). Across two experiments, TE was higher under conditions of more frequent (p=.3) compared to rare (p=.05) probabilities of a task-switch, suggesting that more volatile environments induce less regularity in the order of responses. Increases in TE co-occurred with an increased number of cross-task behaviours, and these responses appeared to be influenced by reduced sensitivity to the odds of success if sticking with behaviours relevant to the more probable task. There is an apparent upside to such altered sensitivity, as those who showed poorer routine maintenance were subsequently faster to adapt to novel tasks that required combining subsets of behaviours from across tasks. Thus, reduced awareness of the odds of success when sticking with a routine may result in increased expectations regarding the relevance of cross-task behaviours to the current situation. Maintaining a routine may therefore be in part dependent on increased expectations of success when sticking with a task, which facilitates exclusion of other task influences.

In this study, we have addressed the challenge that is quantifying the degree to which a routine has been maintained. Our proposed *transition entropy*measure quantifies the extent to which the same order is followed, while being invariant to the number of behaviours that have been used in the ordering; a dissociation not achieved by previous work (Garner et al., 2024; Machado, 1993; Schneider & Logan, 2006; Watanabe et al., 2023). Across two experiments, TE scores were robustly indicative of regular orderings of behaviours, and higher TE scores corresponded to greater flexibility in the face of novel tasks. Thus, TE is a suitable measure for quantifying regularity in the ordering of behaviours, under conditions that require repeated performance of multiple behaviours. The ability to assess self-chosen orderings is novel, as previous investigations have largely relied on fixed “correct” orders (e.g. Carlson & Sohn, 2000; Garr & Delamater, 2019; Graybiel, 1998; Humphreys et al., 2000). Future work should determine the robustness and reliability of the TE measure across other routine scenarios, both in the lab and outside, as well as the cognitive factors that drive variation in TE scores. In addition, ecological validity could be assessed by determining the relationship between TE scores and questionnaire measures of daily routine tendencies (e.g. Ersche et al., 2017).

We have shown across two experiments that more frequent task-switching disrupts routine maintenance. Frequent switching also selectively increases the proportion of cross-task behaviours (Barnes et al., 2026; Garner et al., 2024). Thus, frequent switching appears to perturb routine maintenance by making cross-task behaviours competitive for selection—at least, when there exist no explicit cues to instruct which task *should* be performed. This finding broadens previous investigations into habit formation, which have largely considered the reinforcement mechanisms that drive the concatenation of stimulus-response associations into a sequence (e.g Desrochers et al., 2010, 2015; Dezfouli & Balleine, 2012; Dezfouli et al., 2014). Although such models can account for a range of behavioural phenomena when animals and humans repeatedly perform stimulus-response associations in the lab (see Miller et al., 2019; Watson & de Wit, 2018, for reviews), the resulting frameworks often fail to account for failures in real-world routine maintenance (Gardner et al., 2024). The current findings show that at least part of this gap may be informed by elucidating the mechanisms that determine control over routine performance in uncertain environments.

Having determined that frequent switching perturbs routines via cross-task interference, it is pertinent to ask why cross-task behaviours become more competitive for selection when routine maintenance lapses. By calculating the ongoing odds that the last task remains relevant (the task odds), along with the odds of success given the strategy of sticking with the last relevant task (the success odds), we were able to investigate whether sensitivity to these odds were altered during disrupted routine maintenance. Across both experiments, we observed that frequent switching increased cross-task selections when success odds were lower compared to higher. Note that even if the odds of success is lower, those odds should favour pursuing the last relevant task until those locations have been exhausted, for as long as it is understood that the probability of a task-switch is less than 50%. However, short runs of more frequent switches would push the probability of a task-switch closer to 50%, and consequently the success odds would only be mildly in favour of sticking with the current task, compared to runs with fewer switches. This suggests that recent experiences of frequent switching induces a behavioural strategy geared towards anticipating the next task switch, rather than finding the target in the fewest moves possible. Such behaviour could reflect a task-substitution fallacy, where experiencing more switches promotes the notion that it is adaptive to predict the determinants of switches, rather than sticking with the original task. Alternately, it could be that more frequent switching induced a perception that switch-probabilities exceeded 50%, and success odds were interpreted as favouring pursuit of the other task. Asking participants to provide their estimates of switch probabilities at the end of training would be one means of pinning down which of these explanations better accounts for disruptions to routine maintenance.

It is worth noting that although frequent switching altered sensitivity to varying odds of success over both experiments, the observed strength and direction of this relationship is influenced by the other regressors in the model—namely the task odds and number of switches—which are themselves functionally related to the success odds. Thus, it cannot be concluded from the current analysis that frequent switching uniquely disrupts estimates of success odds to impact routine maintenance. Indeed, the finding that cross-task selections occur more frequently when the success odds are lower implies that cross-task selections increase after periods where the probability of a task-switch has been estimated as higher than average, as these are the conditions that yield a lower estimate of success odds for that trial. Fully dissociating the impact of success odds from task odds would require systematic factorial manipulations, e.g. low and high task switch probabilities could be crossed with low and high numbers of task relevant locations to achieve orthogonal manipulations of both factors. This remains a challenge for the future. However, what the current analysis does show is that frequent switching induces a behavioural strategy that deviates from normative behaviour in a manner that covaries with changes in the unique variance carried by the success odds. This implies a disruption to the psychological representations that support routine maintenance under conditions of uncertainty.

Although future work should seek to pin down the exact causal pathway underlying the current findings, the notion that uncertain conditions can produce counterproductive exploration is not without precedent. When tasked with predicting binary outcomes, people are more likely to match their response proportions to outcome proportions (probability matching, see Myers, 2014; Newell & Schulze, 2016; Vulkan, 2000, for reviews). In line with the notion of the task-substitution fallacy proposed here, such behaviour has been attributed in part to a misaligned goal of predicting sequences, rather than tolerating small proportions of losses to maximise trial-by-trial outcomes (Gao & Corter, 2015). Alternately (but not mutually exclusive), cross-task behaviours could also reflect exploration geared towards increasing utility (e.g. Aston-Jones & Cohen, 2005); finding the target in a cross-task location (against the odds) would allow skipping of at least one task routine. Thus, routine maintenance may be hindered when there is a perceived advantage to performing other tasks—in a bid to either learn about causes of disruption, or to make early gains that shortcut the routine.

A further finding was that frequent switching resulted in more cross-task selections on average, compared to rare switching, beyond what was accounted for by the odds of success, the task odds, the experienced number of switches, and the switch rate. Future work should determine whether this variance can be accounted for by extending the modelling approach to better account for individual differences (e.g. in learning rate), or whether other factors are required to account for disruptions to routine maintenance. It has been shown previously that the conditions of initial task exposure determine what is learned over the course of performance; spatial regularities that are not explicitly cued—such as when a visual target reappears in a particular location of a search array over trials—are learned when the regularities occur from the outset, but not when target locations are randomised for early trials (Jungé et al., 2007). This suggests that people initially adopt a way of performing the task that is not updated when conditions change without a sufficient lack of cost (e.g. Irons & Leber, 2018). Similarly, participants who experienced more early switches by chance may have performed more cross-task selections without updating the rate over the course of the rehearsal phase. Future work should systematically manipulate early switching experiences in uncertain environments to determine if this is a cause of ongoing failures in routine maintenance.

Having identified that frequent switching disrupts routine maintenance by altering sensitivity to likely success when pursuing a task, we sought to explore the cognitive consequences of maintaining a routine. We found that stronger routine maintenance corresponded to poorer performance on a novel task that requires the mixing of routines between tasks, relative to a task that requires replication of a known routine. Note that with the current design we are unable to disentangle whether or not this relationship is dependent on the group difference in routine maintenance. Rather, TE scores may reflect a more sensitive index of the grouping, resulting in significant correlation. Future work needs to determine whether the observed correlation would arise in the absence of a group-based manipulation. Further, although the anticipated relationship between routine strength and transfer bias was observed, the relationship was statistically significant for only one measure (accuracy), and not for a second measure (estimated learning onset). This may be due to reliability differences between the two measures of transfer performance. However, this possibility needs to be verified by future work. Further, this result should be replicated to verify it’s robustness, given that the observed correlation was exploratory. However, this finding tentatively suggests a selective pressure to disruptions in routine maintenance; relative to consistently followed routines, a consequence of less regular ordered responding may be an increased propensity to combine components of learned tasks to solve novel problems. Further work is required to determine whether this increased propensity is due to sensitivity to failure, as is suggested here, or other cognitive processes, such as associative links in memory between task-relevant and cross-task behaviours (cf. Lee et al., 2022).

Here, we have proposed a framework for the theoretical and experimental analysis of the self-ordering of task-relevant behaviours. By determining the transition entropy through a task state space, we can quantify how regularly self-chosen orderings have been followed. This allows us to theoretically define routine maintenance as the reliability with which transitions are made between regularly visited states. This opens a door to the experimental interrogation of routine maintenance, that goes beyond imposing a “correct” order to the routine. We have also shed some insight into why routines may be hard to maintain; uncertain and unstable environments may result in oversensitivity to a lack of reward early in the routine, or a trialling of behaviours from other tasks in a bid to gain early wins. These findings demonstrate that our understanding of why routines are hard to maintain requires consideration of the dynamic and uncertain environments in which they are implemented. Further interrogation of these insights should enable better support of routine maintenance, thereby increasing our understanding of their apparent benefits for daily functioning.

## Method

A detailed description of the experimental method is reported in Barnes et al. (2026). Here we recap the key details for the current study. Barnes et al. (2026) manipulated switching during rehearsal over two experiments, and the consequent impacts on learning transfer in one experiment (Experiment One). Here we analyse the impact of switching on routine maintenance using the data from Experiment One, and we test for replication of effects using the data from Experiment Two. We determine the impact of routine maintenance on learning transfer using the data from Experiment One.

### Participants

All participants were recruited from the paid and undergraduate pools at the University of New South Wales. The study was approved by the UNSW Human Research Ethics Committee (approval number 6341). All participants provided written consent before the session began. *Experiment One* 100 participants were included in the study (M age = 20.3, ±4.49, four unreported; 23 male, 77 female). *Experiment Two* 100 participants were included (M age = 19.3, ±2.66, four unreported, 30 male, 66 female, four use another term).

### Procedure

Experiment One and Experiment Two both consisted of a task acquisition phase and a rehearsal phase. Experiment One also included a learning transfer test.

#### Task Acquisition Phase

The goal of the task acquisition phase was to ensure that participants could perform both tasks with high accuracy prior to the rehearsal phase. Participants completed five practice trials where the grid of squares was presented with a dark grey border and the location of the target was drawn randomly from the 16 squares. The trial began with presentation of the grid of squares, presented in the default grey surrounded by the border. If a non-target square was selected then the square turned dark grey for the duration of the selection before reverting to the default colour. If the target square was selected then the target was presented for a total of 500 ms, including the duration for which the mouse button was depressed. The next trial began once the square reverted back to default grey.

After the five practice trials, participants were told that they would now see the squares surrounded by a coloured border, and that the colour signalled a specific house. Participants were also informed that the animal targets would only hide in four rooms in any given house, and that their task was to learn to find the target in four moves or fewer. They were also informed that if they found the target in four moves or fewer, then a tone would be played that signalled their success. Tones were taken from a 2000 ms piano arpeggio where the number of tones played mapped inversely to the number of moves taken to find the target (e.g. a target found in one move elicited four tones, a target found in four moves elicited one tone).

Participants then performed the task with the first coloured border, which was drawn pseudorandomly from the four colours listed above. The four target locations were pseudorandomly selected across participants, with the constraint that one target location fell in each quadrant of the grid, with one and only one target location in a corner and one and only one target location in the inner quadrant of the grid (see Fig 1B for examples). Participants completed Task *A* for an initial 50 trials and then continued until they met the criteria of finding the target in four moves or fewer for at least 9 of the preceding 10 trials (so a perfect performer could complete the acquisition within 50 trials), or until they completed 200 trials. On completion of Task *A*, participants were instructed that they would now see a new coloured border which signalled that the animals had moved to a new house, and that these animals would be found in four new locations (Task *B*). Task *B* locations were pseudorandomly chosen according to the same criteria as for Task *A*, with the additional constraint that no locations could overlap with those selected for Task *A* (see Fig 1B). Participants completed Task *B* according to the same accuracy criteria as set for Task *A*. They were then instructed that they would complete each task once more, and completed a block of 40 trials in Task *A*, followed by a block of 40 trials in Task *B*. Participants were excluded if they took more than four moves to find the target on more than 20% of trials, for either of the latter Task *A* or *B* blocks.

#### Rehearsal Phase

Note that all participants completed 160 trials of Task *A* and 160 trials of Task *B*. To balance explicit feedback between the two groups, a proportion of trials were randomly allocated as feedback trials. Participants were informed that now they would hear the performance-contingent tones on only a subset of trials, and that the tone signalled how many points they had won. Points were converted to a cash bonus at the end of the session (10-20AUD). Only Stay (non-switch) trials were selected as reward trials, as this guarantees points under the optimal strategy. Further, the number of trials randomly allocated as reward trials was equivalent to 50% of the Stay trials experienced by the Variable group, thus equating the number of possible reward trials between the two groups.

#### Learning Transfer Test

After rehearsal, participants from Experiment One completed two blocks of 40 transfer trials. A new coloured border was used for each block (randomly selected from the remaining two possible border colours). Participants were informed that the new coloured borders represented moving to a new house, and that they may be able to use their previous experiences to learn the new hiding places.

For one block of trials, the target locations were the same as either Task *A* or Task *B*, counterbalanced across participants (identity transfer). For the other block of trials, the target locations were pseudorandomly drawn from both Task *A* and Task *B*, using two squares from each (mixed transfer, see Fig 6). Whether the identity or mixed transfer test was presented first was counterbalanced across participants.

**Figure 6.**
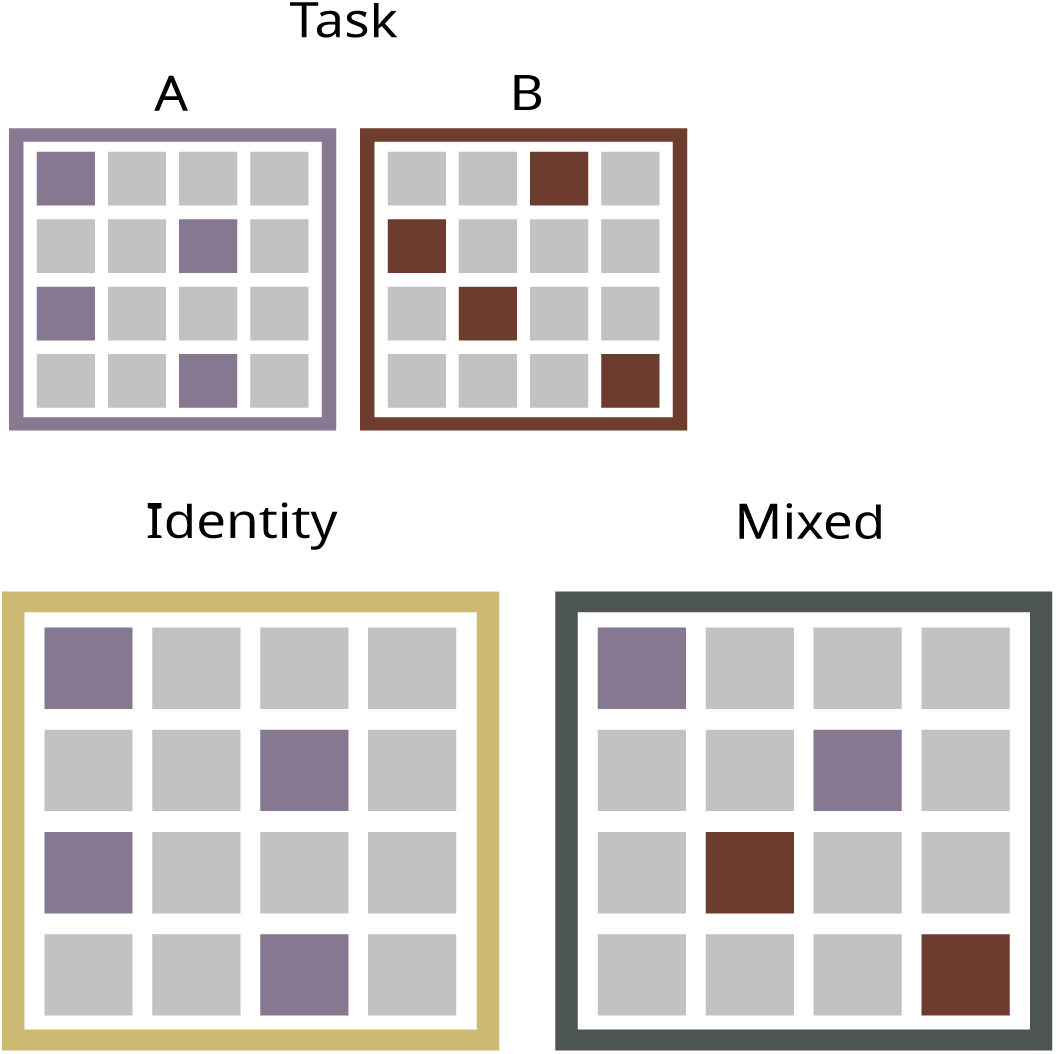
Experiment 1 Test Phase: Two learning transfer tasks were generated using either a complete mapping of either Task A or Task B to a new coloured border (Identity), or a combination of two task-relevant locations from each of Task A and Task B (Mixed).

**Figure 7.**
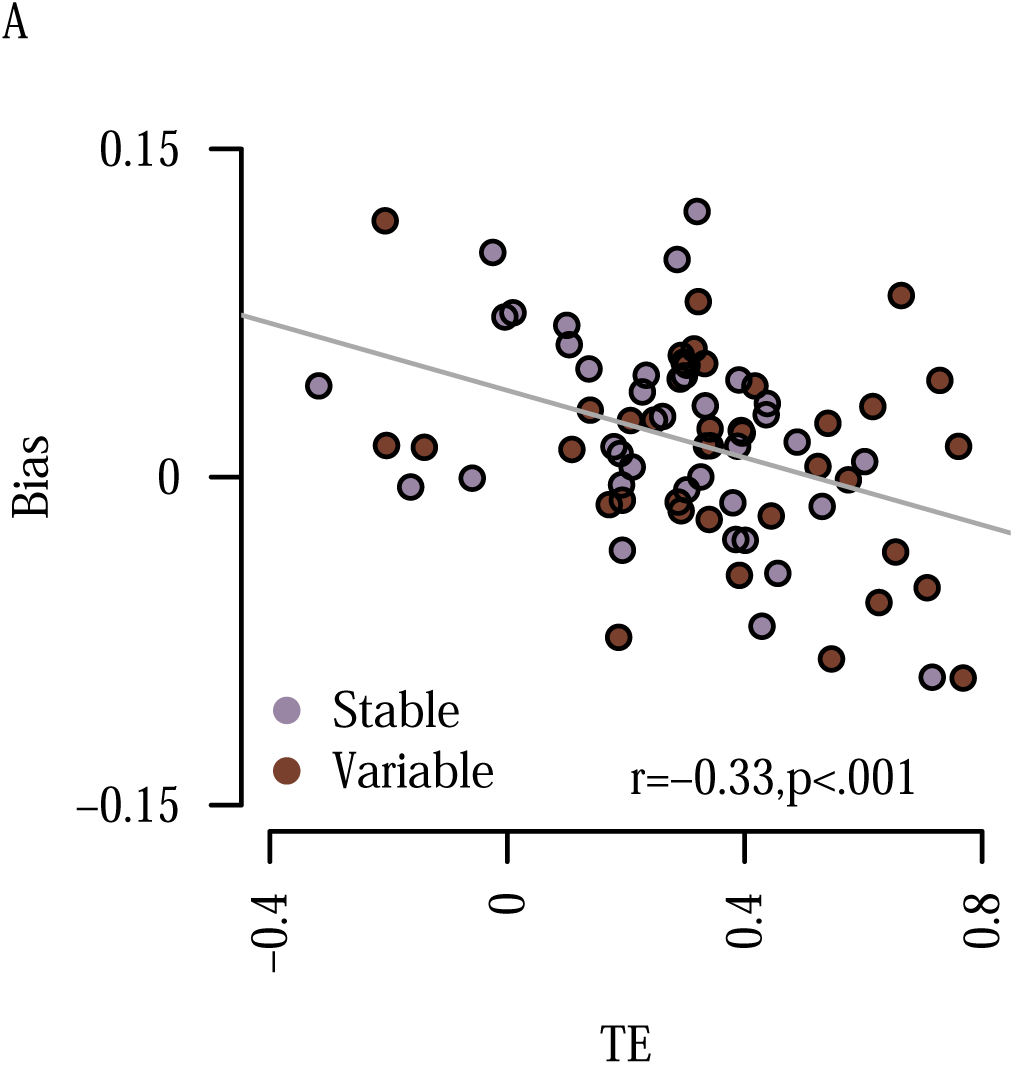
Correlation between transfer bias scores and log transition entropy (TE) scores. Each dot represents one participant.

### Statistical Design

The data is available from the Open Science Framework, and was preprocessed and analysed using bespoke scripts written in R. Preprocessing scripts are available here and the data analysis can be reproduced using the code available here. For the learning transfer analysis, skewed distributions were log transformed, and outliers less or greater than 1.5 times the inter-quartile range (IQR) were removed. This resulted in the removal of 11 participants from the analysis of the learning transfer data.

### Impact of Cross-Task Interference on Routine Maintenance

After determining the impact of switch rate on TE scores, we sought to identify whether the Variable group were more likely to make pre-emptive visits to locations relevant for the less likely task (Barnes et al., 2026) because they formed altered expectations regarding the likelihood of cross-task events. Note that in the rehearsal phase, explicit cues were removed, meaning that participants had to determine which task was most likely to be currently relevant. The optimal strategy under these conditions—for both group Stable and group Variable—was to identify from which task (*A* or *B*) the last target (LT) was found (Task*_LT_*), and to exhaust locations from that task before moving to locations relevant for the task where the last target was not found (Task*_¬LT_*). This can be demonstrated by the following formulation. First, it is necessary to compute the prior probability that the task currently being performed (*T_c_*) is the same task as Task*_LT_* . Specifically, when *n* locations from Task*_LT_*, and *m* locations from Task*_¬LT_* have already been selected and found not to contain a target, the probability that Task*_LT_* is still the current task can be calculated using:

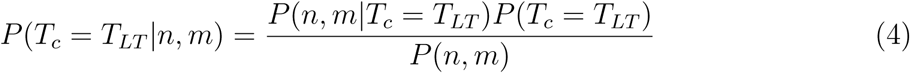

Note that this probability reduces over the course of a trial, as *n* and *m* selections are made. Further, the probability that the task state has changed (to the one for which the last target was not found) can be obtained using:

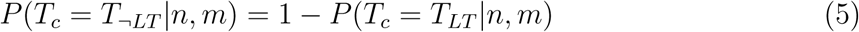

Also note that given equations 4 and 5, it is possible to obtain the odds ratio that quantifies how much more likely it is that Task*_LT_*is still the relevant task, relative to Task*_¬LT_* :

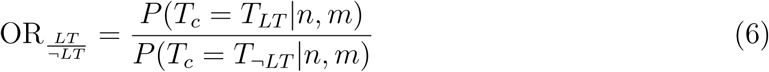

Having calculated the prior probability that Task*_c_* is the same as Task*_LT_*(equation 4), the second step to determining the optimal strategy is to calculate the probability of finding the target, given the remaining locations left to try from each task:

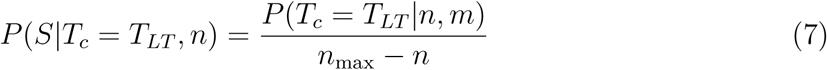

where *S* represents success (finding the target), and *n_max_* is the total number of target-relevant locations in Task*_LT_* (where in this case *n_max_*= 4). Conversely, it is also possible to calculate the probability of finding the target if selecting locations relevant to *T_¬LT_*:

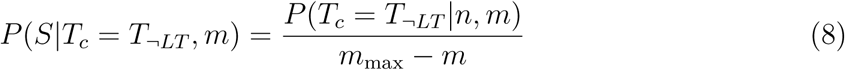

Last, the optimal strategy can be determined by calculating the odds ratio of success if sticking with locations relevant to *T_LT_* :

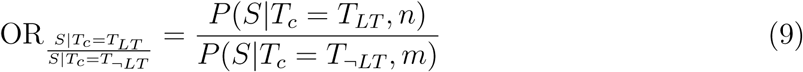

Note that for as long as the true probability of a switch is less than p=.5, and neither *n* nor *m* locations have been exhausted, then 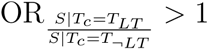. Thus, the optimal behaviour is to exhaust locations from Task*_LT_* rather than pre-emptively switching to locations for Task*_¬LT_*. Further, although the prior probability that Task*_c_* is the same as Task*_LT_* reduces over the course of a trial as *n* and *m* selections are made (equation 4), in this particular case, because the target appears at all task relevant locations with equal probability, 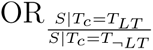 is a constant until all *n* from Task*_LT_* are exhausted.

To determine the extent to which pre-emptive visits to locations from the less likely task were attributable to variation in the odds that the current task was still Task*_LT_* (task odds, equation 6), or the odds of success when performing the remainder of Task*_LT_* (success odds, equation 9), we calculated each odds on a trial by trial basis, dependent on the experiences of each participant. We used a prior task-switch probability of p=.5, that was updated with each experienced switch. We used this prior as participants had experienced equal numbers of trials in each task up to this point of the experiment. This value quickly converged to the correct switch probability. The resulting values were entered into a hierarchical logistic regression where the dependent variable was whether a given location selection was from Task*_LT_* (0) or from Task*_¬LT_* (1). We entered group, task odds, success odds, and their two- and three-way interactions as predictors in the model. We also controlled for general experience of task-switches, by computing a regressor that counted the cumulative number of switches (Sw = Ʃ*_t_*_=1_*^T^S*(*t*), where *S*(*t*) = 1 if the task switched on trial *t*). Note that this predictor would expect a linear increase in Task*_¬LT_* selections over the course of the Rehearsal Phase, as might be expected if switching between tasks caused cumulating confusion. We kept random effects maximal (Barr et al., 2013) by including subject slopes for task odds, success odds and number of switches along with subject level intercepts. Effects indicated as significant from the ANOVA test on model parameters were verified using a simulated likelihood ratio test (Hartig et al., 2026, LRT,): data were simulated using a model that dropped the effect of interest (M0) over 500 iterations. The full (M1) and reduced (M0) models were fit to the simulated data and the difference in log likelihoods was computed. For each simulated LRT, we report the observed log likelihood difference, the mean and standard deviation of the log likelihood differences from the generated null, and the probability of the observed difference. This provides a null distribution of log likelihood differences against which to evaluate the probability of the observed log likelihood difference between M1 and M0. This procedure is preferred to the ANOVA procedure as the latter may produce unreliable statistics given its reliance on the model degrees of freedom which are not clearly defined for mixed models. Note that continuous predictors were trimmed for each participant to remove extreme values (less or greater than 3 standard deviations from the mean), and were downsampled to ten equally spaced segments per group (this ensured that each segment contained multiple behavioural observations). Predictors were then mean centred and entered into the regression model.

## Statements and Declarations

### Availability of data and material

The data is available here here [from the Open Science Framework](https://osf.io/4ehry/overview?view_only=2fba551e23794d78ad1bc4bca583add9).

### Code availability

The code used to run the experiment is available [here from Zenedo](https://zenodo.org/records/16623763). The data were preprocessed and analysed using bespoke scripts written in R which are available [here from Zenedo](https://zenodo.org/records/18385834). The data analysis reported in this paper can be reproduced using the code that can be followed [here on this notebook on GitHub](https://garner-code.github.io/routines_produce-results/).

### Authors’ contributions

KGG & CRN conceptualised the analysis, completed the analysis, produced figures and wrote the manuscript. MLP provided feedback on conceptualisation and analysis, and edited the manuscript.

### Ethics approval

The study was approved by the UNSW Human Research Ethics Committee (approval number 6341)

### Consent to participate

All participants provided written consent before the session began.

### Consent for publication

N/A

